# Causal role for pulvinar burst firing in thalamo-cortical attention control

**DOI:** 10.1101/2025.09.16.676591

**Authors:** Rober Boshra, Michael Harris, Kacie Dougherty, Melissa Berg, Britney M. Morea, Henry J. Alitto, Mario C. Rodriguez, W. Martin Usrey, Sabine Kastner

## Abstract

The primate pulvinar has been implicated in coordinating cortical networks during attention, but its causal role remains elusive. Like other thalamic nuclei, pulvinar neurons switch between tonic and burst firing, but whether bursts influence behavior and cognitive processes is unknown. Here, we show that pulvinar bursts are not only modulated by attention but also predict behavioral performance. In macaques performing a covert attention task, bursts were more frequent for non-cued targets and uniquely reconfigured population codes in parietal cortex. Crucially, electrical microstimulation of pulvinar triggered bursts that enhanced target detection and synchronized cortical spiking, with behavioral benefits scaling with the strength of burst recruitment. These results establish pulvinar bursts as a causal driver of attention, revealing a thalamo-cortical mechanism by which thalamus exerts moment-to-moment control over cortical processing and behavior.

## Main Text

Pulvinar is the largest nucleus in the primate thalamus, and its expansion during evolution scales with that of the neocortex (*1–4*). Unlike first-order thalamic nuclei, which relay information from the sensory periphery to the cortex, pulvinar is a higher-order nucleus that is densely and reciprocally interconnected with cortex (*5*, *6*), placing it in a strategic position to regulate information flow across cortical networks. Empirical evidence in monkeys suggests that pulvinar synchronizes cortical areas during attentional processing (*7*). However, a causal role for pulvinar in driving cognitive cortical networks and influencing behavior has not yet been established in the primate brain (*5*, *6*, *8*).

A distinct feature of thalamic neurons is their ability to alternate between tonic- and burst-firing modes (*9*, *10*). Burst firing (BF), generated by calcium currents following hyperpolarization, is especially prevalent in higher-order thalamic nuclei such as the pulvinar (*11–13*). In first-order thalamus, BF has been proposed to act as a “wake-up call” that rapidly recruits cortex to unexpected stimuli and is followed by tonic firing for high-fidelity relay of information (*9*, *10*, *14*, *15*). Consistent with the “wake-up call” hypothesis, bursts drive cortical responses with greater efficacy and temporal precision than tonic spikes (*14*, *16*). While this relay-centric idea of the thalamus may be applicable to first-order nuclei, it is not clear whether it extends to higher-order thalamus. More fundamentally, how and whether thalamic BF plays functional roles in cognition and behavior in the primate brain has never been empirically tested.

Here, we address these gaps by probing the role of BF in pulvinar and its impact on cortical activity and behavior. We conducted simultaneous recordings from pulvinar and the lateral intraparietal area (LIP), a core node in the cortical attention network that exhibits strong interactions with pulvinar during attention behaviors (*17*, *18*). We examined whether pulvinar bursts occur during engagement in spatial attention and whether they reshape cortical population responses. To establish causality, we combined this approach with electrical microstimulation to probe whether inducing bursts in pulvinar can influence cortical processing and attention behaviors.

### Pulvinar bursting is a functional firing mode during attentional engagement

First, we sought to understand the role of thalamic BF during attentional engagement. We hypothesized that if pulvinar BF indeed plays a role in cognitive function, it would be modulated during an attention task. We probed this idea in two rhesus macaques performing an Egly-Driver task (*19*) in which a spatial cue indicated the location of a target presented at perceptual threshold with high probability (occurrences at cued location: 80%, at non-cued locations: 20%; see Materials and Methods; Fig. 1A). Targets were presented after a variable delay period during which attention was allocated at the cued location and the animals anticipated the onset of the target. After target presentation, animals released a lever to receive juice reward for all correct detections at cued and non-cued locations. During electrophysiology sessions, the two animals detected 79.6% (SD=5.6) of targets at the cued location and 67.7% (SD=9) at non-cued locations, indicating that they used cue information successfully to direct attention to locations of highest relevance for behavior (p<0.001; see Fig. 1B).

**Figure 1:**
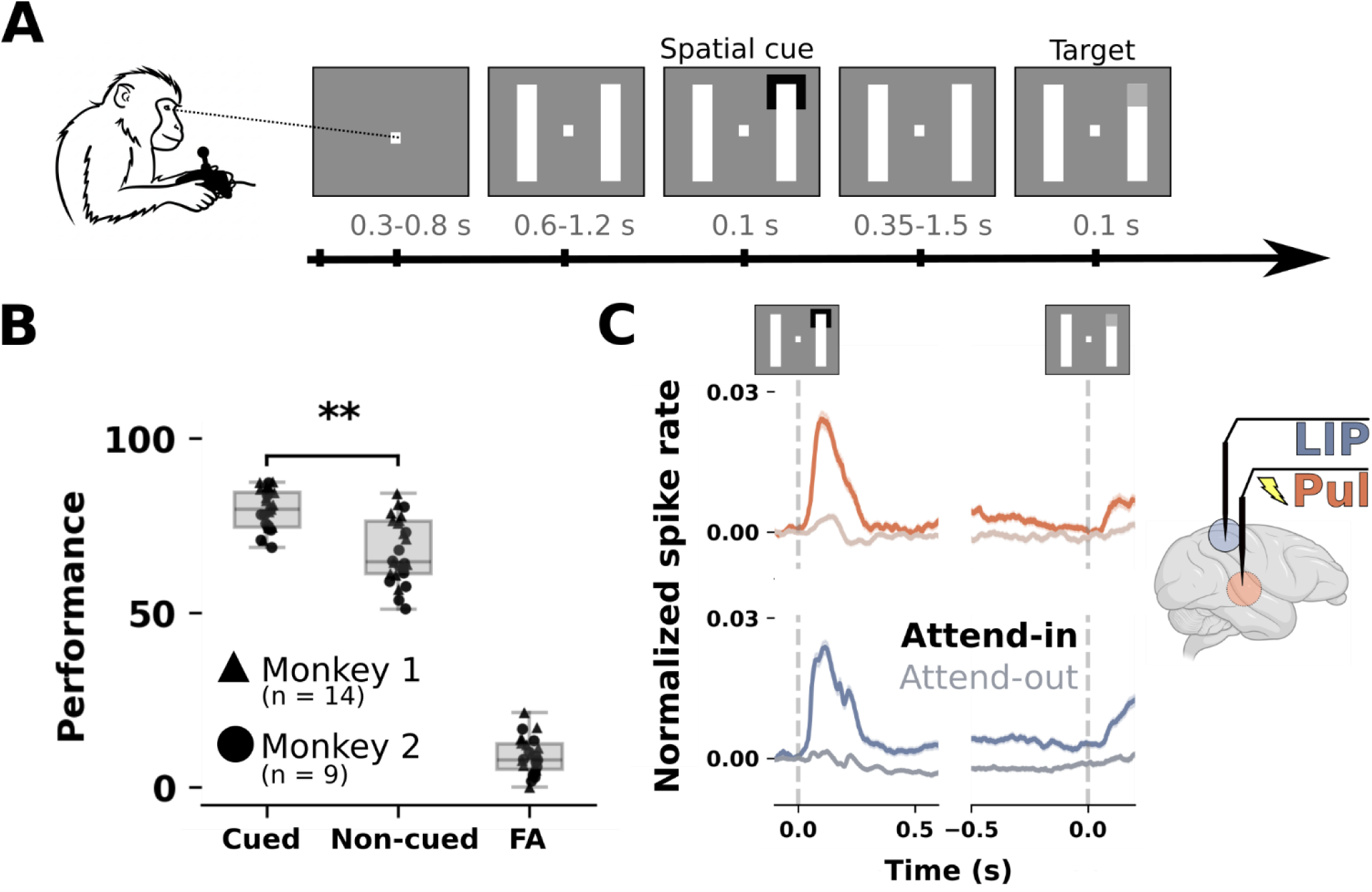
Experimental design, behavioral performance, and recording setup. A) Two rhesus macaques performed an Egly-Driver task using a lever to initiate trials and respond to targets. Central fixation was maintained throughout. B) Both animals performed the task at perceptual threshold, exhibited strong attention effects (i.e. hit rates at cued vs. non-cued locations), and had low false alarm rates. Each session is denoted as a triangle (monkey 1) or circle (monkey 2). Horizontal lines in boxplot denote medians. Box edges are 25th and 75th quartiles. Whiskers indicate the most extreme data points within the 1.5 interquartile range. C) Smoothed and normalized population spike rates for the attend-in (dark color) and attend-out (light color) calculated from cue-responsive neurons in pulvinar (orange) and LIP (blue). Cue-locked and target-locked waveforms are shown on the left and middle panels, respectively. Inset: illustration of simultaneous recording and electrical microstimulation in pulvinar, along with simultaneous recordings from LIP. **: p < 0.001. FA: false alarms.

We acquired simultaneous spiking activity and local field potential recordings from two interconnected areas, dorsolateral pulvinar and LIP (see Fig. 1C, S1; see Materials and Methods). Both pulvinar and LIP exhibited attentional modulation during the delay period when the animals were covertly allocating attention at the cued location (Fig. 1C; Fig. S2A), similar to previous work (*7*, *17*, *20*). Next, we quantified BF in isolated neuron activity as the average number of bursts per trial and neuron. A burst was defined as two or more spikes with inter-spike intervals (ISI) <4ms preceded by a minimum of 100ms of no spiking activity (*21*) (see Materials and Methods). During fixation periods, we found that BF in pulvinar neurons was significantly reduced compared to idle periods during which the animals were not engaged with the task (p<0.01; Fig. S2B, top right), consistent with previous work showing increased BF during idle or rest periods (*22*). However, in contrast to LIP, pulvinar bursting activity increased to levels similar to idle periods after cue presentation (p<0.001, fixation vs. cue; Fig. S2B). These results suggest that pulvinar bursting is a functional firing mode that is not only observable in wakefulness but is also directly affected by task engagement.

Next, we analyzed whether attention modulated the prevalence of bursting in cue-responsive neurons (pulvinar: 617, LIP: 726) during a pure cognitive state, i.e., the delay period (0.25s after the cue offset and until target onset; see Materials and Methods). We found that both pulvinar (p=0.012; average modulation index [MI]=0.01) and LIP (p=0.002; average MI=0.034) neurons had a small but significant increase in BF when the receptive field (RF) of a neuron overlapped with the attended location (attend-in) as compared to when attention was allocated elsewhere (attend-out; Fig. 2A), indicating that attentional engagement does not only modulate cortical bursting, as previously shown (*23–26*), but also thalamic bursting. Critically, our findings show that pulvinar BF is functionally linked to sustained attention even in the absence of external stimuli, revealing that higher-order thalamic bursts contribute to endogenous attentional processes and are not confined to bottom-up ‘wake-up call’ mechanisms.

**Figure 2:**
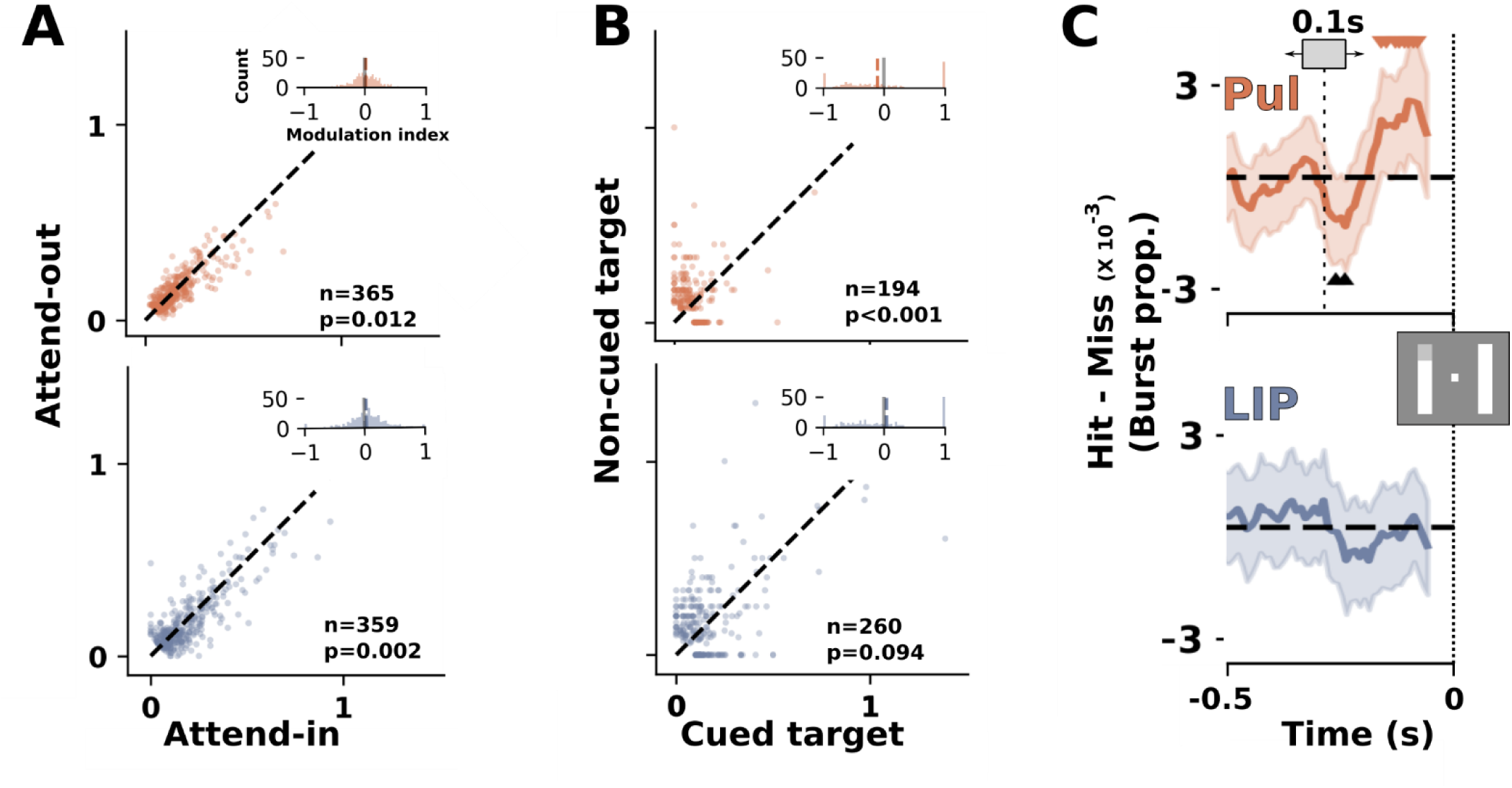
Pulvinar exhibits burst activity during active task engagement. A) Both pulvinar (top) and LIP (bottom) neurons exhibited enhanced burst proportions during the delay period when the spatial cue overlapped with their RFs (attend-in) compared to when the cue was outside their RF (attend-out). Insets show computed modulation indices for bursting. N denotes the number of neurons that had more than 5% burst proportions across the two conditions and were thus included in the analysis (see Materials and Methods). Dashed orange and blue vertical lines on insets denote mean modulation index for pulvinar and LIP, respectively. B) Pulvinar (top), but not LIP (bottom), neurons exhibited a strong increase in burst proportions following the onset of detected infrequent targets at non-cued locations compared to the more frequent targets at cued locations. C) The time-course of the difference between hit and miss trials in bursts across isolated neurons in the delay period for pulvinar (top) and LIP (bottom). The signal is smoothed using a sliding window (schematic shown as grey rectangle) of 100ms and a stride of 10ms where the x-axis denotes the middle of the sliding window. Shaded regions denote the bootstrapped 95% confidence intervals (n=10,000 permutations). Downward carets denote significant windows where bursting in Hit > Miss trials. Black upward carets denote the opposite.

We then investigated whether bursts are related to target selection by contrasting burst proportions for trials when a target was cued (i.e., statistically frequent) versus when it was non-cued (i.e., statistically infrequent). Burst proportions were significantly enhanced in pulvinar for non-cued as compared to cued targets (p=0.001; Fig. 2B, top; average MI=-0.109), suggesting that BF may facilitate the detection of infrequent targets (*9*, *15*). Comparatively, LIP neurons did not exhibit this effect (p=0.094; Fig. 2B, bottom; average MI=0.042). While previous work has proposed that first-order thalamic bursting may be engaged when thalamus neurons process a novel or unexpected stimulus after long quiescent periods (>5s) (*9*), our results suggest that, especially in higher-order thalamus, bursting may occur at much shorter timescales and may have a role in target selection.

Next, we asked whether bursts were linked to behavior by examining trials when animals made a correct detection (hit) versus when they did not report a target (miss). We calculated the smoothed number of bursts, normalized by the number of trials, over time using 100ms windows (sliding by 10ms) before presentation of cued targets (see Materials and Methods). We then utilized a non-parametric bootstrap (n=10,000 permutations) test to extract the time-delineated burst occurrence in the delay period and found that in pulvinar, but not in LIP (Fig. 2C, bottom), BF increased 100ms prior to target onset during a correct detection as compared to a miss (Fig. 2C, top). Conversely, BF in miss trials was enhanced 200ms prior to target onset in pulvinar (Fig. 2C, top). Together, these results suggest a temporally-specific link between thalamic BF and target perception.

### Pulvinar bursts boost cortical population-level representation of attended location

Pulvinar influences on cortical sensory processing (*7*, *27*) and cognitive control areas (*17*, *18*, *28*) have been well-established. However, the mechanisms by which pulvinar affects cortical processing remain elusive. Our demonstration of attentional modulation of pulvinar bursting activity related to behavioral performance suggests BF as a possible signaling mechanism to influence cortex and to aid in guiding attentional allocation. Based on computational modeling work (*29*) proposing a putative role for pulvinar signals in sustained cortical firing (e.g., sustaining spatial location information during spatial attention or working memory tasks; Fig. S2), we hypothesized that moment-to-moment spiking of pulvinar neurons may bias the population-level representation of the attended location at the cortical level towards the visual space represented by these pulvinar neurons. Specifically, we expected that, as bursts have higher efficacy of eliciting cortical responses (*14*, *30*), they would impose a stronger bias in the LIP-represented attended location as compared to tonic spikes. To probe this hypothesis, we trained regularized logistic regression models to decode the moment-to-moment attended location from cortical spiking populations (Fig. 3A; see training and axis-extraction details in Materials and Methods). We trained the models on binned spiking (isolated and multi-unit) activity during correct, cued-target trials to distinguish between attended ipsilateral and contralateral locations (2 configurations; see Fig. 3A).

**Figure 3:**
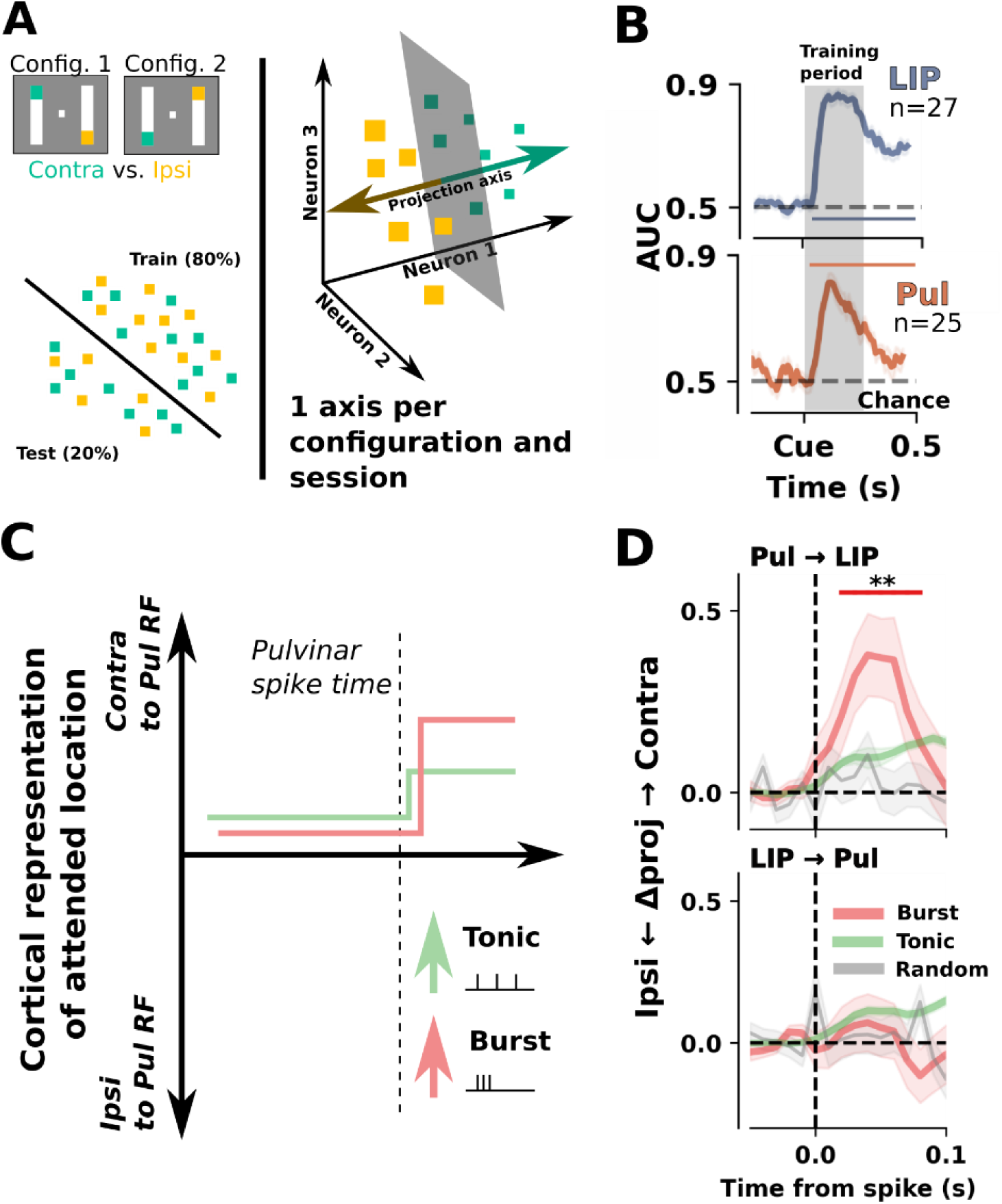
Pulvinar bursts bias cortical population-level representations towards the attended location. A) We trained logistic classifiers to decode the attended location (binary: contralateral vs. ipsilateral; 2 configurations) using population activity in response to the cue. To examine the trajectory of the population’s encoded attended location representation over time, we extracted a decoder’s classification hyperplane and projected the data on the attention location axis (contralateral vs. ipsilateral; right). B) Aggregated model performance on unseen test data scored using area under the curve (AUC) of the receiver-operating characteristic curve for LIP (top) and pulvinar (bottom). Shaded grey rectangle denotes the period used for training (performance on training data not shown). Shaded colored regions denote standard error of the mean. C) Schematic of our hypothesis that a pulvinar burst would bias the decoded attended location in cortex to the same RF of a pulvinar bursting neurons. We hypothesized that a tonic spike would have a smaller but similar effect on cortex. D) Directed influence of one area on another’s population-encoded attended location. Top: Pulvinar bursts were linked to a significant bias towards contralateral space. Tonic spikes induced a trending, but much smaller effect in the same direction. Randomly sampled (grey) times decoded using the same method showed no effects to serve as control. Bottom: Neither LIP bursts nor tonic LIP spikes induced a significant bias in pulvinar-encoded attended location. Shaded regions denote the standard error of the mean. **: p<0.001.

Models for both LIP and pulvinar populations were able to decode the attended (i.e. cued) location in unseen test data (20%), extending into the delay period after cue offset (Fig. 3B). To examine the effect of a source area’s spiking on a target area, we extracted the contralateral-ipsilateral encoding axes and projected onto these axes the target area’s population spiking data, time-locked to spiking activity in the source area during the cue-target delay period. Finally, we calculated the normalized change in encoded attended location (in the target area) to a 50 ms window preceding spiking activity in the source area. We found that pulvinar tonic spikes induced a small trending effect of a shift in encoded attention location towards contralateral space representation (overlapping with the RF of pulvinar spiking; p=0.13). This effect was four times larger when the pulvinar spiking event was a cardinal (i.e., first) spike of a burst as compared to a tonic spike (p<0.001; Fig. 3D, top). In contrast, LIP spiking did not influence the representation of attended space in the pulvinar population activity (Fig. 3D, bottom). Taken together, these spike-population coupling results suggest that pulvinar spiking activity biases cortical representations towards the attended location. Specifically, thalamic bursting may constitute a powerful thalamo-cortical mechanism to boost maintenance or guide an attentional shift to the respective visual field location (*29*, *31*, *32*).

### Pulvinar electrical microstimulation induces a neural cascade to influence cortex

Thus far, our results indicate that pulvinar BF is modulated by attention, linked to behavior, and influences cortical processing during sustained attention. Next, we sought to causally probe the influences of thalamic BF on cortical processing and behavioral performance. We electrically microstimulated (EMS) pulvinar using biphasic 16-pulse trains delivered at 200 Hz prior to target onset in the Egly-Driver task (Fig. 4A; see Materials and Methods). EMS enables temporally accurate perturbation of a local neuronal population; thus, we delivered EMS in one of four staggered time windows prior to target onset to assess temporal specificity of any causal effects. To better understand local responses evoked by our EMS protocol, we simultaneously stimulated and recorded from pulvinar (see Materials and Methods; Fig. S4). Following EMS, we found a robust neural cascade that started with a brief increase in firing, followed by global inhibition, and ended in vigorous pulvinar firing (Fig. 4B, in contrast with a gradual return to baseline firing rates in cortex post-EMS (*33*)). This cascade was observable at the single-trial level in the raw data (Fig. 4B, left) as well as in isolated single neurons (Fig. 4B, right).

**Figure 4:**
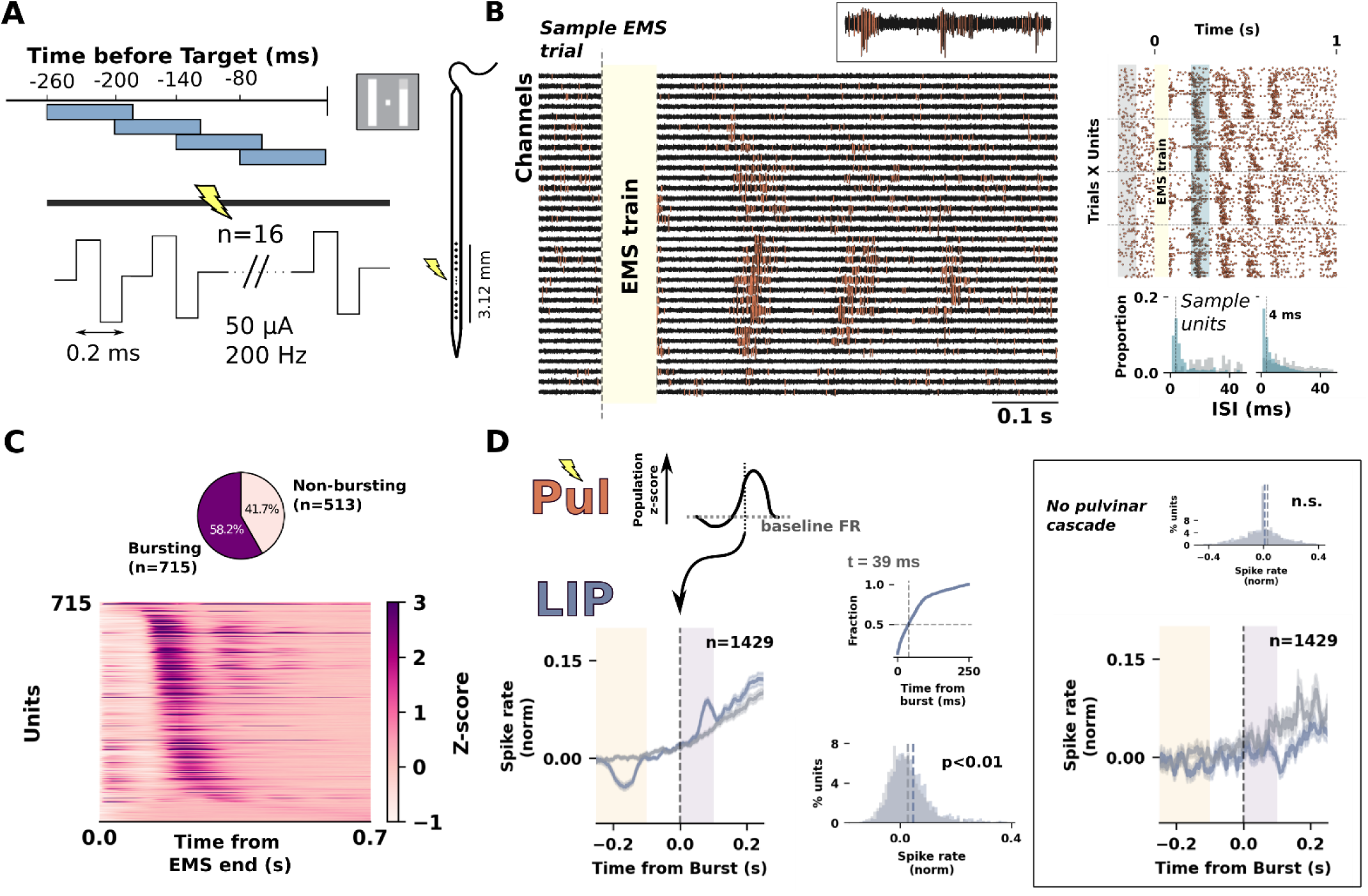
Pulvinar electrical microstimulation elicits bursting and affects cortical activity in LIP. A) In 50% of trials, we delivered biphasic pulse trains (n=16; 200Hz; 0.2ms) to pulvinar in one of 4 windows prior to target onset. To examine the effect of EMS, we simultaneously stimulated and recorded from pulvinar (see Materials and Methods). B) Left: sample highpass-filtered data showing that following EMS (yellow rectangle), local pulvinar activity (32 channels) exhibited a robust response cascade starting with brief excitation, extended inhibition, and a rebound in activity with pronounced increased firing that could repeat. Orange waveforms denote spike threshold crossings. Inset: zoomed-in view for a single channel. Right, top: raster of isolated neurons in a sample session and their response to EMS in 4 sample trials. Right, bottom: ISIs shifted to predominantly fast spiking (<4 ms) following stimulation (cyan) compared to baseline (grey). C) Top: The majority of pulvinar neurons exhibited significant increases in spiking activity, denoting a burst response following EMS and the ensuing inhibition period (58.2%; see Materials and Methods). Bottom: normalized firing rates for the neurons exhibiting post-EMS bursting ordered by their firing peak times. Note the subsequent firing peaks in a large proportion of neurons following the primary peak. D) Pulvinar bursting was coupled with increased firing in LIP neurons and multi-units. Left, top: Pulvinar burst time was estimated as the crossing of the combined normalized firing rate of all bursting neurons (panel C) of a 0.675 threshold (see Materials and Methods). Left, bottom: smoothed spike rates for LIP units time-locked to pulvinar threshold crossings for EMS (blue) and sham (grey) trials. Shaded regions denote the standard error of the mean. Top, right: LIP exhibited increased spiking at 39ms that was significantly higher than time-matched sham trials (Bottom right). Inset: In instances when EMS did not elicit a burst, no cortical responses were observed. Conventions follow outside of the inset.

Critically, the rebound from the inhibitory period resulted in characteristic BF, such that the neurons’ ISIs exhibited a strong shift to under 4ms (Fig. 4B, lower right). We also observed repetitions of this cascade such that some neurons returned to an inhibition period followed by a second vigorous firing period (see Fig. 4B and C). This periodic response suggests that stimulation altered the underlying dynamic trajectory of pulvinar neurons (*12*), potentially engaging excitatory-inhibitory loops (e.g. thalamocortical collaterals in TRN (*34–36*)).

Out of the total 1,228 isolated neurons, 715 (58.2%) showed significant post-EMS firing rate (FR) enhancement indicative of burst activity (see Materials and Methods). To visualize this responsive population, we normalized (z-scored) spiking responses in EMS trials to the baseline period for each neuron and ordered pulvinar neurons according to the peak of their normalized FRs after EMS train offset (Fig. 4C, bottom). The repeated neural cascade was observable across a large proportion of neurons with a distinct inhibition (shown in white) prior to the bursting rebound. Together, these results indicate that EMS elicited a robust and characteristic response in the local pulvinar population.

Next, we examined the effects that pulvinar EMS-evoked bursting had on LIP neurons. We predicted that, as thalamic bursts may have higher efficacy in eliciting a response in interconnected cortical regions (*10*, *14*), LIP spiking activity would be coupled to burst times (also see (*35*)). We estimated the population-level burst time on a single-trial level by taking the first time point at which the population of stimulation-responsive neurons (see Fig. 4C) exceeded a firing threshold. To combine across neurons, we averaged the z-scored FRs and selected the first timepoint that passed a threshold of z=0.675 (corresponding to an alpha of 0.1). Next, we computed LIP population spiking time-courses, time-locked to the estimated pulvinar population-level burst times. We found a significant increase in FR following pulvinar burst time in LIP compared to time-matched sham trials (50% of significant population peaking at 43 ms after pulvinar estimated burst onset; see Fig. 4D). To rule out the possibility that the cortical firing was artifactual or simply coupled to the stimulation train and not the elicited bursts, we analyzed EMS trials that did not surpass the z-score threshold compared to pre-stimulation as indicative of successful delivery of current but no successful elicitation of bursts. These trials showed no increase in cortical spiking (Fig. 4D, inset), suggesting that stimulation-evoked BF in pulvinar indeed led to an increase of spiking activity in LIP neurons.

We then sought to understand the extent of spatial selectivity given that pulvinar neurons with specific RFs were stimulated by limiting our analysis to LIP neurons that were exclusively responsive to only one of the two locations in contralateral visual space (i.e., upper versus lower visual field locations in the same hemifield; see Fig. 1A). We found that LIP neurons exhibited a post-burst increase in FR only when their RF overlapped with that of the stimulated pulvinar site (p<0.001; Fig. S3). Our results show that EMS-elicited bursting had similar characteristics as compared to intrinsic bursting. That is, BF was able to recruit the interconnected LIP neuron populations during active task engagement in a spatially selective manner.

### Pulvinar electrical microstimulation increases target detection

We previously established that intrinsic pulvinar BF was linked to attention behavior (Fig. 2C). Therefore, we next examined whether we could probe this relationship causally using EMS-induced thalamic bursting by calculating hit rates (HR) for EMS and sham trials. As pulvinar neurons in one session predominantly represented contralateral RFs within a quadrant, we split trials during which cues directed attention to the contralateral visual field (i.e., overlapping with the stimulated pulvinar site) and to the ipsilateral visual field (i.e., non-overlapping and opposite to the stimulated hemisphere). We then calculated the difference in HR between EMS and sham trials for both ipsilaterally and contralaterally cued trials for each session. Given the delayed onset of bursts relative to EMS, we split our analysis between trials with EMS onsets that were more than 150ms before target onset (early EMS; sufficient time for pre-target burst activity to occur) and trials where EMS onsets were less than 150ms (late EMS; burst activity occurs following target presentation). We hypothesized that EMS would enhance detection of contralateral, but not ipsilateral targets, when EMS-elicited bursts preceded the target (see Fig. 2C).

Strikingly, our results showed that stimulation significantly increased HR in the contralateral condition (Fig. 5A, left; mean=4.9%, p=0.01). This effect was not observed for either the ipsilateral condition (Fig. 5A) or when the EMS onset did not provide enough time for pre-target bursting (p>0.05; Fig. 5A, inset), indicating that pulvinar EMS had both temporally and spatially specific effects on behavior and causally enhanced target detection in the visual field corresponding to the stimulated region in pulvinar. To rule out that EMS influenced behavior through recruitment of motor circuitry or induction of phosphenes (*37*, *38*), we analyzed eye movements and false alarms in EMS trials. EMS did not exhibit an increase in false alarm rates, did not alter pupil dilation metrics, and did not influence eye movements (Fig. S5), suggesting that our EMS behavioral effects were purely cognitive. These results provide causal evidence to support the temporally-specific manner by which pulvinar BF is linked to behavioral performance (see Fig. 2C).

**Figure 5:**
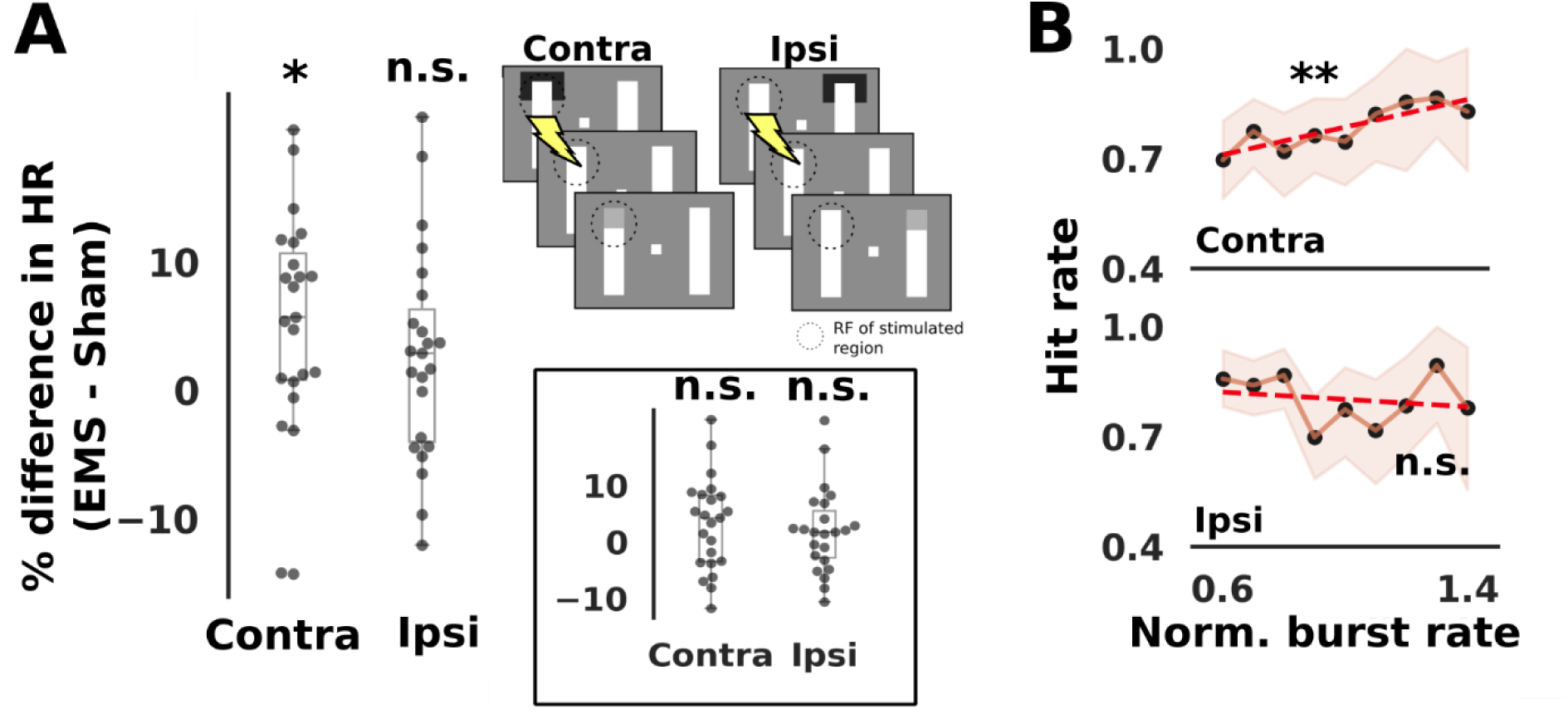
Pulvinar EMS recruits bursting mechanisms to improve target detection in a spatially specific manner. A) Compared to sham, EMS increased hit rate for targets overlapping with the pulvinar site’s RFs (p=0.012), but not when the targets were presented ipsilateral to the pulvinar EMS site (p>0.05). This boost in performance manifests contralaterally for trials where EMS onset was delivered at least 150ms prior to target presentation, but not when stimulation was delivered after that (inset). B) Hit rate in EMS trials was linked with pulvinar BF. Increased BF corresponded to higher HR when EMS was applied contralateral to the target (top; p=0.005) but not when the target was presented ipsilaterally to the target (bottom; p>0.05).

Our previous results support a role for pulvinar bursting in attention and demonstrate that pulvinar BF increases spiking activity in LIP (Fig. 2, 4D). Thus, we examined whether EMS-induced pulvinar bursts can be directly linked to changes in behavioral performance (see Materials and Methods). To probe this, we computed a bootstrapped (n=10,000) HR distribution as a function of the population-level normalized pulvinar spiking activity (0.1-0.2 s post-stimulation; stimulation-responsive neurons only; see Fig. 4C).

Results show a linear positive correlation between the pulvinar burst response and HR (Fig. 5B, top; red regression line, r=0.83, p=0.005). As expected, this effect was not observed in the ipsilateral condition (Fig. 5B, bottom; p>0.05). These results indicate that the effects of EMS on behavior are driven by the bursting component of the ensuing cascade and establish a causal link between thalamic bursting and attention behaviors.

## Discussion

Thalamic BF has been mechanistically well characterized. BF is driven by calcium currents that follow the release from inhibitory input, which hyperpolarizes thalamic cells and de-inactivates T-type Ca^++^ channels. This de-inactivation allows for subsequent depolarizing inputs to generate a large Ca^++^ potential upon which rides a rapid train of Na^+^ based action potentials—the *burst* (*39*). Here, we show that this mechanism is not only able to powerfully influence cortical processing, but can also closely be linked to behavioral performance, thereby suggesting a critical function of thalamic BF in moment-to-moment attention control. Critically, we provide evidence not only for a correlative relationship of thalamic BF to attention behaviors but also establish a causal relationship by inducing bursts through EMS. These results indicate for the first time a role for higher-order thalamic BF in cognition and behavior. At the same time, they establish a causal mechanism for pulvinar in controlling attention behaviors.

A remarkable feature of T-type Ca²⁺ channels is that their gating, and the BF they enable, can be recruited across remarkably different brain states and processing conditions for distinctly different purposes. During sleep, prolonged inhibitory input hyperpolarizes thalamic relay neurons, de-inactivating T-type channels so that rebound depolarization generates rhythmic bursts (*39*, *40*). These bursts drive spindle oscillations and widespread thalamocortical synchrony, effectively disengaging information exchange between thalamus and cortex. In contrast, similar channel dynamics can be transiently recruited during wakefulness to enhance information exchange to unexpected stimuli (*9*). While this role has not been linked to behavior, it has been studied in first-order thalamic nuclei, where non-preferred stimuli induce the inhibition needed to de-inactivate T-type Ca²⁺ channels, thereby allowing BF to occur in response to abrupt transitions to a preferred stimulus. In this context, BF has a very different role, to strongly drive postsynaptic cortical targets and enhance the transmission of sensory information. Thus, the same ionic mechanism underlies both the slow, rhythmic bursting that characterizes non-wake states and the rapid, environment-driven (i.e., exogenously driven) bursting that can sharpen sensory processing during wakefulness. Here, we show for the first time a functional role for thalamic BF that is causally linked to cognitive behavior. We found that BF is recruited during attentional task engagement of higher order thalamus, modulating interconnected cortex on the neuron (Fig. 4D) and population (Fig. 3D) levels.

While theoretical considerations have often linked thalamic BF to attention functions (*9*, *15*, *41*), these ideas have never been tested empirically. Here, we show that in pulvinar, BF contributes to endogenous attentional processes and target selection. Specifically, we show that thalamic BF is linked to target detection in temporally specific ways both for intrinsic bursting as well as EMS-induced bursting, thereby linking BF to behavioral performance. Indeed, the strength of EMS-induced burstiness scaled with behavioral success (Fig. 5C). Further, our results suggest that pulvinar BF supports the maintenance of spatial location information by biasing cortical population responses towards the attended location during delay periods. Previous work has shown that maintenance of spatial location information in cortex can manifest as discrete short events of increased activity (*31*, *32*, *42*, *43*). While these discrete events have been linked to cognitive behavior, they have primarily been attributed to local intra-areal dynamics and inter-areal cortical signaling (*31*, *32*, *44*). However, given that such events appear to be coordinated in time across areas, a common thalamic coordinator of such events may present an alternative mechanism (*7*, *18*, *29*). If so, our results advance the novel hypothesis that pulvinar BF serves as a thalamo-cortical mechanism, which is causally coupled to cortical discrete events and contributes both to their coordination and ongoing maintenance of spatial location information.

Although pulvinar has long been implicated in attention and other cognitive functions based on its rich anatomical connectivity with cortex (*45–48*), empirical evidence for such roles has been sparse. The most compelling evidence for a role of pulvinar in attention control has come from lesion studies in humans and monkeys. Structural or functional pulvinar lesions lead to attention deficits, which can range from profound, as in visuo-spatial hemineglect (*49–51*) (i.e. a failure to direct attention to contralesional space), to mild in the form of slowing of orienting responses to contralesional space (*38*, *52*). Our results shed new light on the neural mechanisms affected in these studies and provide a mechanistic framework for understanding pulvinar and higher-order thalamus that is necessary for developing strategies to improve attention. Specifically, we demonstrate that pulvinar BF acts as a causal driver of attention, recruiting thalamo-cortical circuits to exert rapid, moment-to-moment control over cortical processing and behavior.

## Materials and Methods

### Experimental model and subject details

We collected data from two adult rhesus macaques (*macaca mulatta*; 11 and 9 years old and weighing 15 and 12kg, respectively). All procedures were approved by the Princeton University Animal Care and Use Committee, conforming to the National Institutes of Health guidelines for the humane care and use of laboratory animals.

### Behavioral task and training

We trained the two animals to perform a modified version of the Egly-Driver task (*19*) using operant conditioning. The animals were trained to press and hold a lever to initiate a trial and to release the lever to report detection of a target presented at sensory threshold. We used Presentation software (Neurobehavioral Inc.) to control stimulus presentation, dispensing of reward, utilization of the eye-tracker, and generation of triggers to align electrophysiological recordings to task events. The animals’ gaze was monitored using an eyelink 1000Plus eyetracker (SR Research, Ottawa, ON). Any deviation from 2 degrees of visual angle (dva) surrounding the fixation point aborted an ongoing trial and required the animal to release the lever before proceeding with another trial.

In the Egly-Driver task, each trial was composed of a variable fixation period (0.3-0.8s), after which horizontally- or vertically-oriented bars were presented such that the four ends of the two bars were at 10 dva. After a variable period (0.6-1.2s), an informative visually salient visual cue was presented for 0.1 s at one of the possible target locations. In 80% of trials, the target was presented after a variable delay period (0.35-1.5s) at the cued location (cued trials). In 20% of trials, the target was presented in one of the two equidistant target locations. To encourage the animals to only report true detections, we designated 10-15% trials globally as catch, such that no target was presented and the animals received juice reward for maintaining a lever press throughout. Trial type as well as cue and target locations were randomly set following the designated percentages every trial.

### Neuroimaging data acquisition

Animals were scanned for surgical planning and post-surgical electrode placement confirmations. They were sedated with ketamine (1-10mg/kg i.m.) and dexmedetomidine (0.007-0.02 mg/kg i.m.) before placement in an MR-compatible stereotaxic frame (1530M; David Kopf Instruments, Tujunga CA). Vital signs were monitored with wireless ECG, pulse, respiration sensors (Siemens AG, Berlin), and a fiber optic temperature probe (FOTS100; Biopac Systems Inc, Goleta CA). Body temperature was maintained with blankets and a warm water re-circulating pump (TP600; Stryker Corp, Kalamazoo MI).

We acquired whole-brain structural MRI data collected either on a Siemens 3 Tesla MAGNETOM Skyra or on a Siemens 3 Tesla MAGNETOM Prisma using a Siemens 11-cm loop coil placed over the head. T2-weighted volumes were acquired with a 3D turbo spin echo with variable flip-angle echo trains (3D T2-SPACE) sequence (voxel size: 0.5mm, slice orientation: sagittal, slice thickness: 0.5 mm, field of view (FoV): 128 × 128 mm, FoV phase: 79.7%, repetition time (TR): 3390 ms, echo time (TE): 386 ms (Skyra) or 387 (Prisma), base resolution: 256 × 256, and acquisition time (TA): 17 min 41 s (Skyra) or 15:39 (Prisma). T1-weighted structural images were collected using the 3D Magnetization-Prepared Rapid-Acquisition Gradient Echo (MPRAGE) sequence, voxel size: 0.5 mm, slice orientation: sagittal, slice thickness: 0.5mm, FoV: 128 × 128 mm, FoV phase: 100%, TR: 2700 ms, TE: 3.27 ms (Skyra) or 2.78 ms (Prisma), inversion time (TI): 850 ms (Skyra) or 861 ms (Prisma), base resolution: 256 × 256, and TA: 11 min 31 s.

### Electrophysiological recording and surgical procedures

All surgical procedures were performed under general anesthesia. Animals were sedated with ketamine (1-10mg/kg i.m.) and dexmedetomidine (0.007-0.02 mg/kg i.m.). Animals were then maintained with ketamine and/or isoflurane (maintenance 0.5– 2%) and/or propofol (15-30 mg/kg/hr) under strictly aseptic conditions. We used ceramic cortical screws (Thomas Recording, inc.) and bridging bone cement (Simplex P) to affix a headpost and two customized polyetheretherketone (PEEK) recording chambers (over frontal and parietal cortex) on the animal’s skull (Rogue Research Inc.). The implant parts were custom designed based on MRI-reconstructions of each animal’s skull to tightly fit the contours of the animal’s head and to minimize cement usage (Fig. S1B). Contained within each chamber, we drilled 10-15mm diameter craniotomies using a piezoelectric drill (Mectron Inc.) to provide access to regions of interest. For precise spatial and angular alignment, placement was guided intraoperatively with the stereoscopic Brainsight system (Rogue Research Inc.), calibrated to fiducial markers placed and localized using MR imaging. Custom 3D-printed combined grids and microdrive holders (Fig. S1B) were designed with 3D modeling software based on fused individualized structural scans, implant components, and recording hardware (Autodesk Inventor; Autodesk Inc.). These 3D-printed grids were affixed acutely on cranial chambers using set screws to guide the placement of probes to target regions.

Regions of interest (ROI) were defined on individual animal MRI scans by non-linear warping of the NMTv2 atlas using Advanced Normalization Tools (ANTs; (*53*)) and followed area boundaries defined in (*54*). We recorded from dorsolateral pulvinar, which is anatomically connected to LIP (*55*). For precise estimation of LIP ROI to align with chosen RF eccentricity and angle we also utilized retinotopic maps from (*56*) visualized using AFNI (*57*). Following implantation, we conducted a set of mapping days (3-5 days) to confirm correct placement of linear arrays in our selected ROIs and RFs. Alignment of RFs across recording areas was done using a conjunction of the Egly-Driver task and an RF mapping task where a series of horizontal and vertical bars were flashed to cover the visual space and RFs were estimated using reverse correlation. In cases when we needed to alter trajectory, mapping days were used as needed to supplement in-situ localization scans (see below). Mapping days were not included in data analysis. Across sessions, recorded neurons were predominantly responsive to a single visual quadrant located contralateral to the site of recording (and stimulation in case of pulvinar).

Recordings started at least 2 weeks following craniotomy surgeries to enable post-surgical recovery (*58*). On recording days, we fixed the animal’s head using a metal holder that coupled to the implanted headpost (Rogue Research Inc.). We independently lowered linear probes with microdrives (NAN Instruments). We used Diagnostic Biochips 128-channel deep arrays or Plexon 32-channel probes connected to an Open Ephys acquisition box (monkey 1) or an Intan Recording system (monkeys 1 & 2). Probes were lowered into the brain through sharp custom guide tubes that enabled access through the dura without compromising probe integrity. Signals from probe contacts were digitized at 30 kHz then processed offline to extract spiking activity (see artifact correction and spike-sorting section) and local field potentials (1 kHz sampling rate; 0.1-300 Hz). Each probe was referenced to its guide tube.

In addition to daily tracking of probe depth and its conformance with anatomical landmarks (e.g. brachium of the superior colliculus as a depth limit for pulvinar recordings) and their electrophysiological properties, we localized our target ROIs using two separate MR-guided approaches. Conducted at least twice for each animal, we placed tungsten electrodes in-situ using custom-built 3D-printed MR-safe microdrives and confirmed the location and depth of the electrodes by examining the “shadow” of the electrode tip (see Fig S1A, inset). This was supplemented by trajectory extrapolation scans that were conducted at interim periods between the in-situ scans. In trajectory extrapolation scans, we passed tungsten electrodes through sterilized glass microcapillaries (350 microns inner diameter) to limit electrode bending in the space between the grid hole and the dural surface. The chamber was then filled with diluted betadine (in saline), which provided sufficient contrast for accurate detection of the straight line defining the electrode trajectory in 3D space (using 3D Slicer (*59*)). These lines were extrapolated into the brain to simulate trajectories with respect to target locations and depths.

Recordings presented here were a subset of a multi-component study that leveraged data from the dorsal aspect of inferotemporal cortex, superior colliculus and the frontal eye fields where animals performed other tasks beside the Egly-Driver. These tasks were performed daily after the animal completed >= 800 trials of the main task and will be discussed elsewhere.

### Electrical microstimulation

Trains of current pulses were delivered to target areas of interest using an Intan Stimulation/Recording Controller (Intan Inc.) through a single contact in a 32-channel Plexon probe (Fig. 4A) or through a single tungsten electrode (FHC inc.) placed 1mm away from the pulvinar recording probe (lowered using the same microdrive). During an initial mapping period, a variety of pulse current strengths, lengths, and frequencies were tested, and we chose settings that enabled elicitation of robust local responses in pulvinar (see Fig. 4B) while minimizing total delivered current and delivery time. The following parameters were used for our EMS experiments: 16-biphasic cathode-leading pulses, each with 0.2ms duration, delivered at 200 Hz and with an amplitude of 50 μA. Pulse trains were delivered at one of the following predefined times prior to target onset: 0.26, 0.2, 0.14, or 0.08. We delivered EMS in 50% of all trials. In the case of catch trials, the current was delivered relative to the end time of the trial. EMS was not delivered if the animal’s behavior resulted in termination of a trial (e.g. breaking eye fixation).

For synchronous recording and stimulation, we utilized synchronized Intan controllers. We used an Arduino board customized to generate digital pulses pseudo-randomly every 2-7 seconds. These pulses were routed to the Recording (for LIP) and Recording/Stimulation (for pulvinar) controllers for synchronization using the unique temporal pulse sequence over the course of a recording.

### Artifact correction, signal processing and spike sorting

Strong artifacts were prevalent during the delivery of a pulse train. These artifacts often saturated the amplifier at the time of delivery, especially for pulvinar, and added a slow artifact that manifested as an exponential decay function from delivery time to baseline (see Fig. S4A). Due to saturation, these artifacts were not linearly separable from neural data across channels and were not constant, i.e., the artifact waveform changed dramatically between trains. Accordingly, average template subtraction and other template-driven approaches to artifact removal could not be successfully utilized (*60*, *61*). As artifact removal was necessary for spike-sorting, we employed a custom algorithm shown in Fig. S4B. For every pulse and channel, we performed the following: first, we estimated the 4^th^ order exponential function that matched the period 0.7-4ms following the fast artifact and subtracted it from the original signal. Next, we removed the linear trend from the resultant signal. We then linearly interpolated the fast artifact −0.7-0.7ms locked to pulse onset. To remove signal discontinuities, we applied a DC shift of the post-artifact signal to match the pre-pulse median of 5 samples. We adapted this algorithm as part of spike-interface package’s remove_artifact function (v0.102.0).

Artifact-corrected broadband data were sorted using Kilosort4 (v4.0.21 ^41^). Sorted data were manually curated into isolated neurons and multi-units using the Phy (v2.0b6) graphical user interface. In total, we recorded from 1338 isolated neurons and 1088 multi-units from pulvinar collected across 23 sessions. Identical procedures were followed for LIP data during synchronous multi-area recordings (n=20). In total, we recorded from 1478 isolated neurons and 1674 multi-units in LIP.

We defined a neuron’s receptive field by first normalizing (z-score) each neuron’s firing rate by a random surrogate from the −0.35-0s baseline (n=5000 repetitions). We then calculated a neuron’s normalized response 0.02-0.25s following each of the cue types (4 locations; Fig. 1A). A neuron’s RF was designated as the maximally responsive cue location with significantly increased firing rate for 10 consecutive sampling bins (10 ms). A total of 721 and 726 single neurons were cue-responsive in pulvinar and LIP, respectively. An additional 507 and 703 multi-units were cue-responsive in pulvinar and LIP, respectively.

To visualize smoothed spike rates in response to task events (Fig. S2), we convolved spike times with a causal half-gaussian (sigma=50 ms). To combine neurons with varying firing rates, we normalized each neuron’s activity by its maximum firing rate across all conditions (e.g. attend-in and attend-out). For plotting purposes, each neuron’s normalized firing rate (cue- and target-locked) was baseline-corrected according to the −0.2-0s window before cue onset. To ensure that the target-locked firing activity did not include the transient following the spatial cue, we plotted trials with cue-target delays longer than 0.75s (Fig. 1, S2).

### Behavioral analysis

We assessed the behavioral performance in trials where targets were presented by calculating hit rates (number of hits / total number of trials). We limited hits to when the animals responded with a response time (RT) +/- 3 standard deviations from their mean response time. We utilized performance during catch trials to estimate the rate by which animals responded when a target was not presented (i.e., false alarms). False alarm rates were estimated as 1 - the proportion of correct catch trials (correct rejections). Trials where animals responded prior to cue onset or moved their eyes beyond the fixation boundary were discarded from our analyses.

We calculated the effect of EMS on behavior by computing the percentage differences in hit rates between EMS and sham trials when the target and cue locations matched (cued). We tested this statistically (compared to a null-effect of no EMS difference) by performing a Wilcoxon rank-sum test (two-sided). To test whether the effect of EMS was temporally selective, we split our trials into whether the EMS-target delay was >0.15s to denote whether bursting would occur prior to target presentation. Similarly, we computed this for ipsilaterally cued target trials.

To assess whether the effect of EMS on behavior scaled with burst strength, we binned EMS trials according to their maximal post-EMS FR in 0.1 bins (z-scored; see Population-level neural responses to electrical microstimulation). For each bin, we sampled with replacement from the selected trials and calculated the hit rate. We repeated this process 10,000 times to estimate the 95% confidence intervals (Fig 5C; shaded regions). We conducted this for trials when the cue and target overlapped such that they were either contralateral or ipsilateral to the pulvinar stimulation site. The scaling of HR as a function of pulvinar population-level BF was statistically tested for both conditions using linear regression (linregress function in scipy v1.12.0).

### Burst detection and quantification

A burst was defined as two or more consecutive spikes with an ISI < 4 ms occurring in an isolated neuron and preceded by more than 100ms of no spiking. These are conservative criteria (*22*) that aim to capture action potentials crowning a low-threshold spike, which results from de-inactivated T channels followed by a release from cell membrane hyperpolarization ^14,43^. Although cortical areas have been shown to exhibit BF through different neurophysiological mechanisms that are not exclusive to our thalamic ISI definitions for BF (*23*, *24*, *26*, *62*), we apply the same criteria to LIP neurons for an analogous comparison. We quantified the number of bursts across trials in burst proportions, defined as the average number of bursts elicited by a neuron per trial. For defined time-windows, we calculated burst proportions for a buffer on the two sides of the window to enable accurate detection of bursts, i.e., a cardinal (first) spike in a burst sequence occurring at the beginning of a window.

Although thalamic BF has been reported during wakefulness, previous work has shown it to be attenuated during fixation, especially in first-order thalamic nuclei (*22*). Thus, we estimated the modulation of burst activity by our task structure in all neurons irrespective of cue responsiveness. We calculated burst proportions in five windows: idle (Idle; 0.5s windows with intertrial intervals > 2 s), fixation (Fix; −0.5-0s pre cue periods), cue (Cue; 0-0.5s post cue periods), as well as peri-target presentation periods (pre- and post-target for −0.5-0, and 0-0.5s time-locked to target presentation, respectively).

Neurons with less than 5% burst proportion were excluded from further analyses (Fig. S2B). We computed the bootstrapped differences in pulvinar bursting between fixation and all other windows by sampling 10,000 times with replacement. Differences between windows were then tested using the Wilcoxon rank sum test. To account for multiple comparisons, we used the Holm-Bonferroni method (*63*).

To examine the modulation of burst activity during the delay period, we extracted burst proportions across all trials, accounting for the variable cue-target period, 0.25s post cue offset to time of target onset. We computed burst proportions in all Pul and LIP neurons for trials in which the cue overlapped with a neuron’s RF (attend-in) versus when the cue was outside the neuron’s RF (attend-out). We used a non-parametric bootstrap procedure (n = 10,000) to test for statistical differences between conditions, computing the distribution of differences and estimating a two-sided p-value based on the proportion of resampled differences that crossed zero in either direction. For visualization (Fig. 2A and B, insets), we computed the BF modulation index, which is defined as (*burst_attend−in_* − *burst_attend−out_*) / (*burst_attend−in_* + *burst_attend−out_*). For target selection, we extracted burst proportions in a 0-0.5s post-target window for trials when the target was cued (cued) vs. when the cue did not overlap (non-cued) with the target location. The same procedure and statistical test were conducted as in the delay period.

To probe whether bursts are linked to behavior, we extracted the time of the cardinal spike across all neurons and grouped them in the delay periods of hit and miss trials. We then calculated the smoothed difference between the two conditions over the course of the delay period. For each 100ms window (stride=10ms), we sampled with replacement (n=10,000) from all the bursts across the delay period, then selected burst times within the window and computed the average number of bursts normalized by the total number of hit and miss trials. Lastly, we used the 95% confidence interval (alpha=0.05) to compute the significant time points during the delay period where hit trials had more bursts on average compared to miss trials and vice versa (denoted by carets in Fig. 2C).

The level of a thalamic neuron’s hyperpolarization is known to be coupled to the minimum silent period required to classify a series of fast spikes (ISI <4ms) as a calcium-driven burst (*13*, *21*). As its elicited hyperpolarization voltage is expected to be extreme (see inhibition period, Fig. 4B, left), EMS would shorten time-voltage relationships. Thus, we used high-frequency firing as a surrogate for estimating whether a neuron exhibited burst activity following EMS (Fig. 4C). Based on the response profile of pulvinar neurons to EMS, we defined a pulvinar neuron as EMS-responsive if its post EMS activity (.125-.325s post-train offset) was significantly larger than the pre-EMS baseline (−0.2-0s; Wilcoxon rank-sum test; Fig. 4C).

### Population representation of attended location

To estimate the moment-to-moment changes of encoded allocation of attention in population activity, we trained regularized logistic regression models (scikit-learn^45^ v1.4.2) to classify between contralaterally- and ipsilaterally-cued trials (Fig. 3A). Models that are able to decode between the two conditions form axes in high-dimensional space that denote the extent by which the population activity aligns with one of the two conditions (Fig. 3B; also see (*64*)). As the population responses to a cue are expected to share a common representation with sustained attention (*32*, *65*) (also see Fig. 3B), we trained our models exclusively on the 300 ms following cue onset. For each session, we extracted binned cortical spike counts from both isolated neurons and multi-units in 50ms windows (shifting by 10 ms) spanning a window 0. to 0.3 s following cue onset. The model was trained on 80% of the trials. During the training procedure, model hyperparameters were tuned to optimize for the ability of the model to classify on cross-validated slices of the training data (folds=5). The models were trained using stochastic gradient descent (SGD; SGDClassifier function in scikit-learn) using hinge loss and combined penalty terms (L1 and L2 regularization). To account for the large number of units compared to trials available for each condition, we applied principal component analysis (PCA) following data normalization. Hyperparameters were selected using the grid-search method from the following: i. PCA components to satisfy variance explained: [0.8, 0.85, 0.9, 0.95], ii. Regularization strength (alpha): [1e-5, 1e-4, 1e-3], iii. Ratio of L1 to L2 regularization: [0.5, 0.55, 0.6, 0.65, 0.7], and iv. Learning rate: [1e-5, 1e-4, 1e-3]. This approach was adopted to optimize model performance in a generalizable manner across different sessions and configurations, which may have different ideal parameters depending on data idiosyncrasies. For each session and configuration, the ideal set of hyperparameters was then used to train a final model on the full training set. The performance of these models on cue-locked unseen testing data is shown in Fig. 3B.

Following training, the contralateral vs. ipsilateral encoding axis can then be extracted from the model represented by the formula:

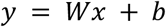

Where y is the predicted class, x is the input feature vector, W is the weight matrix learned through SGD, and b is the learned bias term. Assuming a static threshold at 0, y>0 and y<0 would denote ipsilateral, and contralateral attention, respectively. Note that the input feature vector is normalized and is projected to PC-space. Thus, to project an input spike-count vector v to encoding axis (y^a^) where v is an n-long vector (n units; time dimension is omitted here for clarity), the process is as follows:

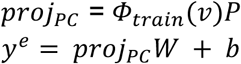

Where Φ_train_(v) = (v – μ_train_) /σ_train_ is the normalization function (z-score) and P is a matrix of dimension *n* × *p*; p is the cross-validation selected number of principal components.

To probe the effect of a source area’s spiking on a target area’s population-level encoding of attended location, we extracted spike counts from the target population in 50ms windows, time-locked to the following events in the 0.5s window prior to target presentation: i. cardinal spike times from bursts in the source area, ii. tonic spike times from the source area, or iii. randomly sampled times with no spike from the source area. Finally, the projected activity was baseline corrected to the 50 ms prior to the time-locking event to assess the time-delineated influence of the source area’s spiking on the target area’s population (Fig. 3D).

### Population-level neural responses to electrical microstimulation

To estimate the trial-by-trial BF timing across pulvinar, we first calculated the normalized single-trial activity for each EMS-responsive neuron, z-scored to the pre-EMS period (−0.2-0s). As different neurons may have varying times of peak activity, we took the timepoint where the average z-score for all EMS-responsive neurons crossed z=0.675 as an estimate of pulvinar burst time.

To investigate the influence of EMS-elicited bursts on LIP activity, we calculated the average smoothed spike rates for all isolated neurons and multi-units that were cue-responsive (i.e., with a RF contralateral to stimulation), time-locked to the estimated time of pulvinar bursting. To estimate the smoothed firing rate for sham trials at a time-course that matches with EMS trials, we randomly sampled EMS-to-target durations from EMS trials and applied these time points and their respective estimated pulvinar burst times to calculate a sham time-course that accounts for neuron-specific statistics. To test whether pulvinar EMS elicited a response in LIP, we compared the smoothed firing rates in the 0.-.1s window following the estimated pulvinar burst time between EMS and sham trials using the Wilcoxon rank-sum test. To test whether LIP activity is strictly coupled to pulvinar BF and not generally to EMS, we instead time-locked LIP smoothed spike rates to the maximal z-score timepoint in EMS trials where the maximum average post-EMS z-score for all EMS-responsive neurons did not surpass 0.675.

To test whether pulvinar’s EMS-elicited bursts to trigger spiking in LIP was spatially selective, we constrained our analysis to neurons that were maximally modulated by one of the two contralateral locations (upper and lower visual fields). We calculated the modulation between the two locations using an index: 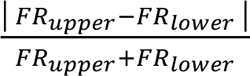 where || denote the absolute value. Neurons with index > 0.95 were selected in this analysis. For each session, neurons were then grouped with respect to whether their RF overlapped or did not overlap with the pulvinar site’s RF for that session (Fig. S3).

## Acknowledgments

The authors thank Michael Arcaro for resources used for receptive field and region of interest localization and Mark Pinsk for assistance with imaging. We thank Kirsten Gerhart, Juan Mercado, and the Princeton Primate Research (PRR) staff for support.

## Author contributions

Conceptualization: RB, KD, SK Methodology: RB, KD, HJA, WMU, SK

Investigation: RB, MH, SK Resources: BMM, MB, MCR Formal analysis: RB Visualization: RB

Funding acquisition: RB, WMU, SK Supervision: WMU, SK

Writing – original draft: RB, WMU, SK

Writing – review & editing: RB, MH, KD, MB, BMM, HJA, MCR, WMU, SK

## Data and code availability

Data supporting all figures will be made available upon publication of the manuscript. Data are available from the corresponding authors upon reasonable request. Code supporting analyses discussed in the manuscript are either from community-supported packages directly referenced in the manuscript or are custom-written. Custom scripts will be made available upon publication of the manuscript.

## Supplementary Materials

Materials and Methods Figs. S1-S5

---

**Supp Fig. 1:**
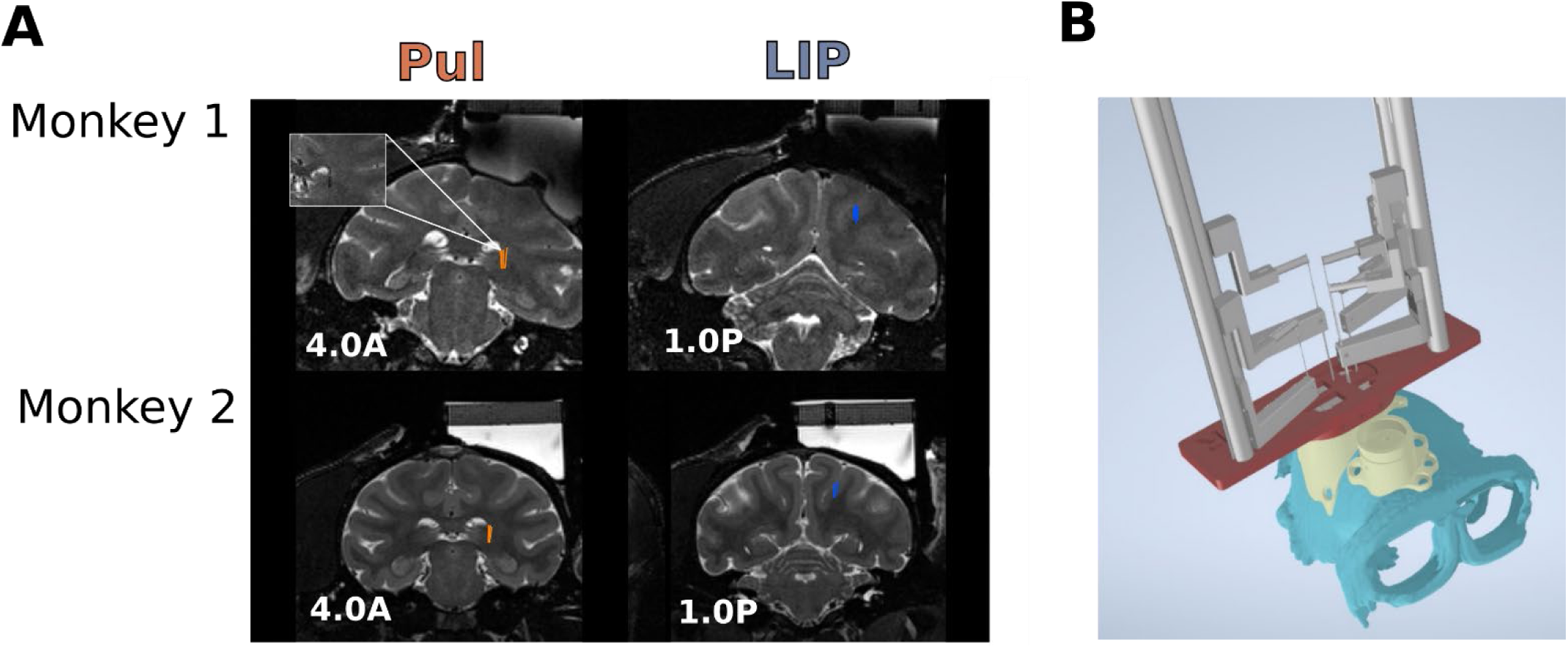
Anatomical locations and localization. A) T2-weighted MRI scans showing the targeted dorsolateral pulvinar (left) and LIP (right) for monkey 1 (top) and 2 (bottom). B) custom fused grids and microdrive holders were designed to give access to our areas of interest, maintaining RF overlap. cyan=reconstructed skull from T1-weighted scan, yellow=custom skull-fitting recording chambers, red=custom 3d-printed grid and microdrive holder, and grey= microdrive towers and probe holders.

**Supp Fig. 2:**
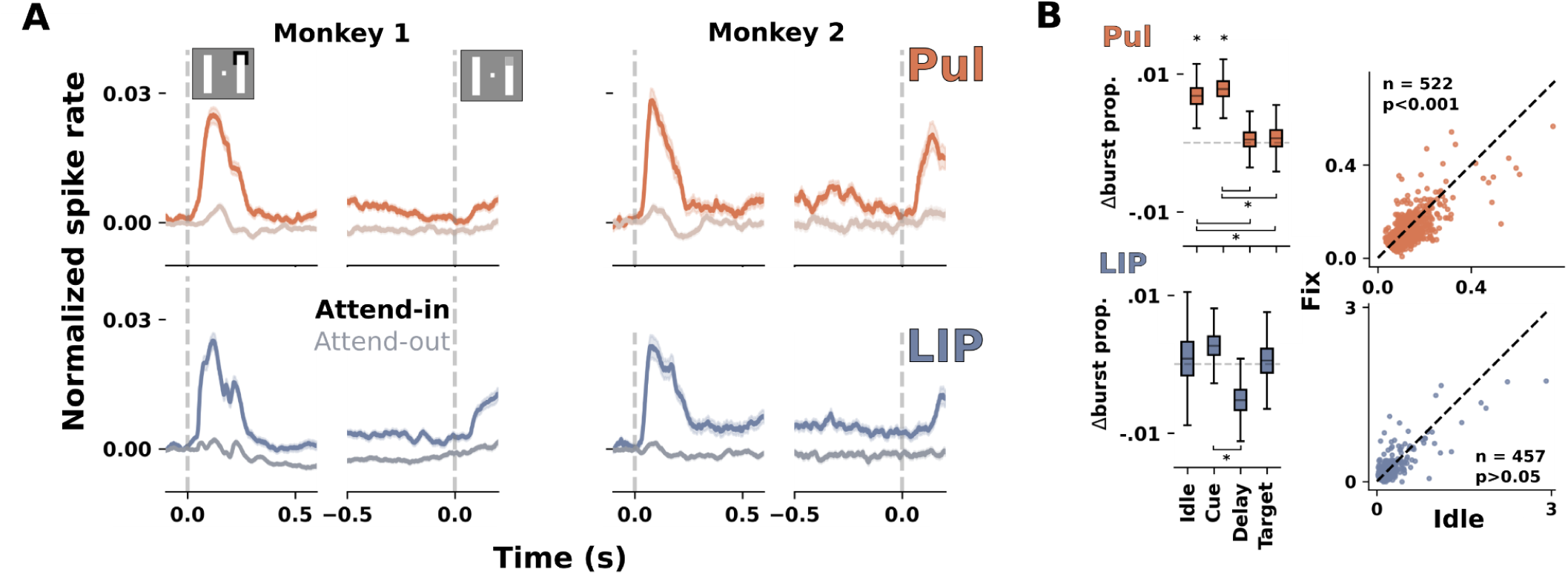
Attentional modulation of population spiking and burst activity in pulvinar and LIP. A) Enhanced sustained activity is observed in peri-stimulus time histograms of the attend-in compared to the attend-out condition across pulvinar (top) and LIP (bottom) for monkeys 1 (left) and 2 (right). Shaded regions denote the standard error of the mean. B) Left: prominence of bursting in pulvinar (top) and LIP (bottom) across different task periods in comparison to the fixation period (−0.5-0 s from cue onset). Pulvinar had elevated bursting during Idle and Cue periods, which were significantly elevated from delay and target periods. Metrics were calculated for neurons that exhibited a mean bursting > 5% of Idle and Fix trials. Right: scatter plot of burst proportions for single neurons in the Fix vs. Idle condition. Pulvinar, but not LIP, had significantly higher bursting during Idle compared to Fix periods.

**Supp Fig. 3:**
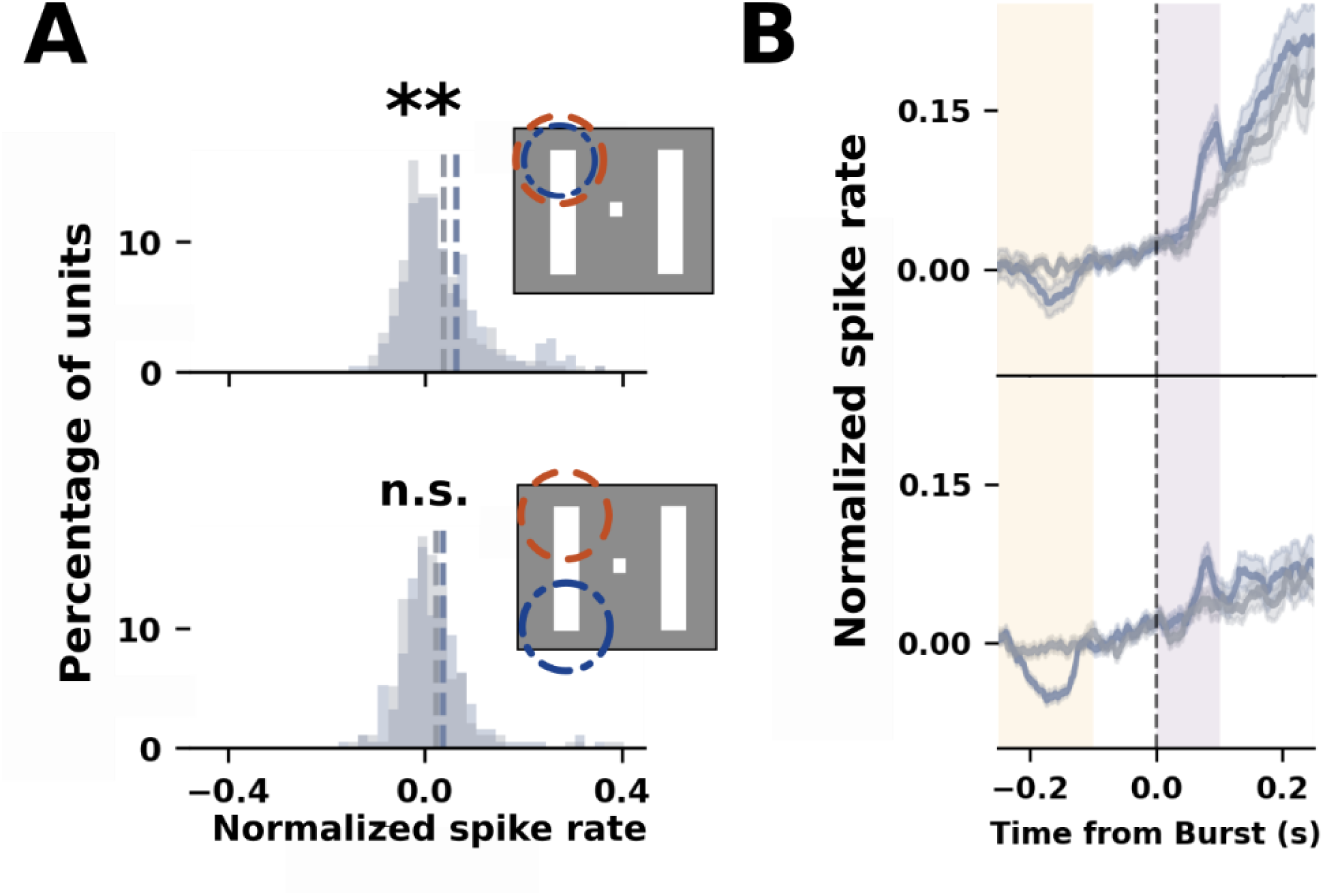
Pulvinar EMS-elicited bursts induce a spatially selective response in cortex. A) Distribution of LIP spike rates in response to EMS-elicited bursts (blue) and sham (grey). LIP isolated neurons and multi-units that shared the pulvinar site’s RF had increased FR following pulvinar bursts (top; p<0.01). LIP units did not exhibit a change in FR when their RFs did not overlap with pulvinar (bottom; same hemifield; p>0.05). Dashed circles denote an illustration of RFs for pulvinar (orange) and LIP (blue). B) time courses for the LIP units with RFs overlapping (top) and not overlapping (bottom) with the pulvinar RF. Conventions follow Fig. 4.

**Supp Fig. 4:**
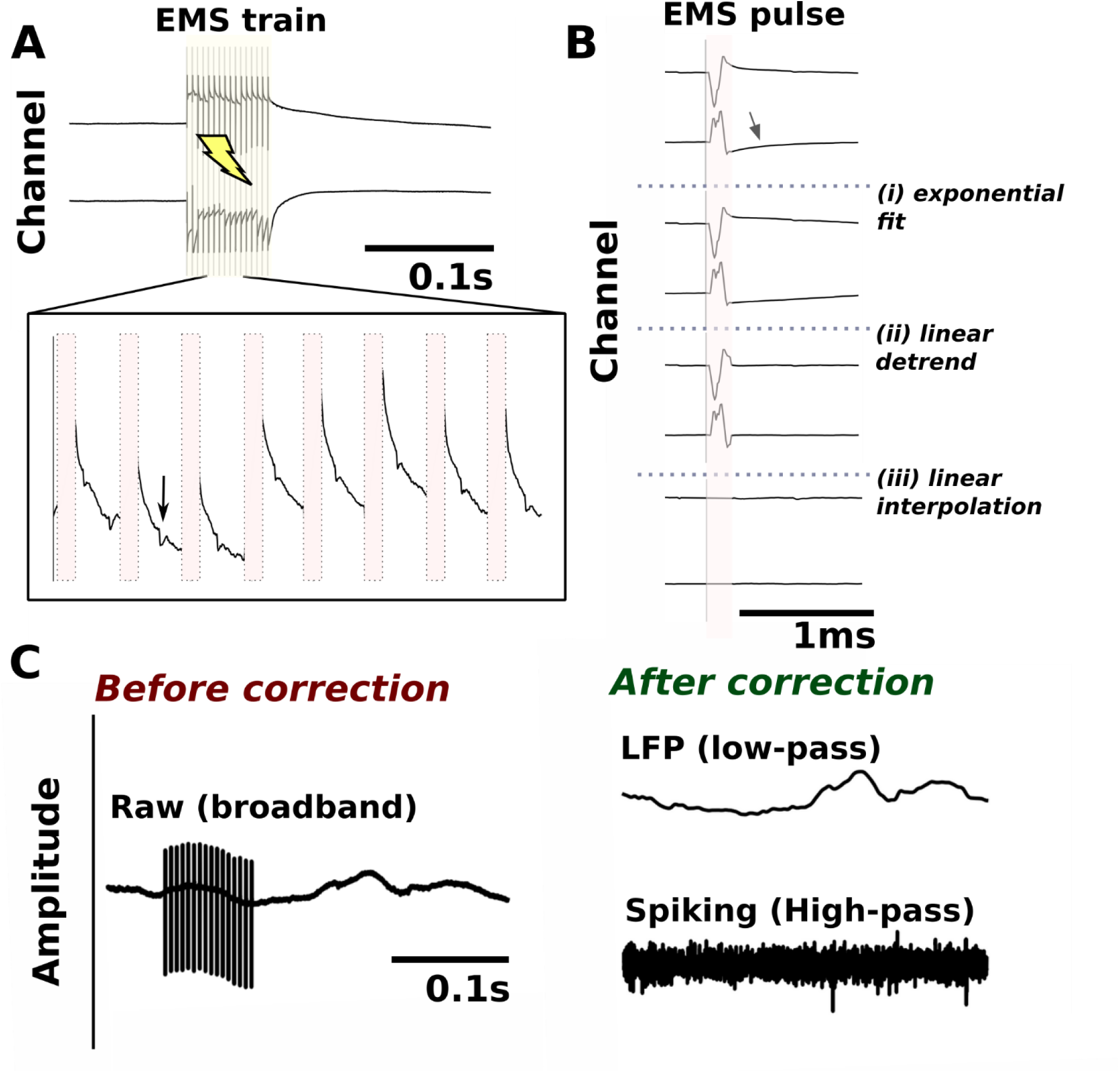
Stimulation artifact patterns and custom correction approach. A) Sample artifacts in two channels (broadband) during EMS in pulvinar. Stimulation often saturated the amplifier and the artifact was not consistent in shape (inset; interpolated pulse duration in red). Neural spiking was often observed between stimulation pulses (inset arrow). B) To reduce the effects of artifacts on our data, we first fit an exponential decay function to model and correct for the slow artifact following the initial pulse and subtracted the fit (i). Then, we computed the linear trend from the end of the pulse to the end of the correction window (ii). Lastly, we excised 2 ms around the duration of the stimulation pulse and linearly interpolated the gap, accounting for any discontinuities due to DC shifts (iii). C) Sample raw data with artifact (left) and the artifact-corrected data of the same snippet following low and high-pass filtering for extraction of local field potentials (top) and spiking (bottom) data, respectively.

**Supp Fig. 5:**
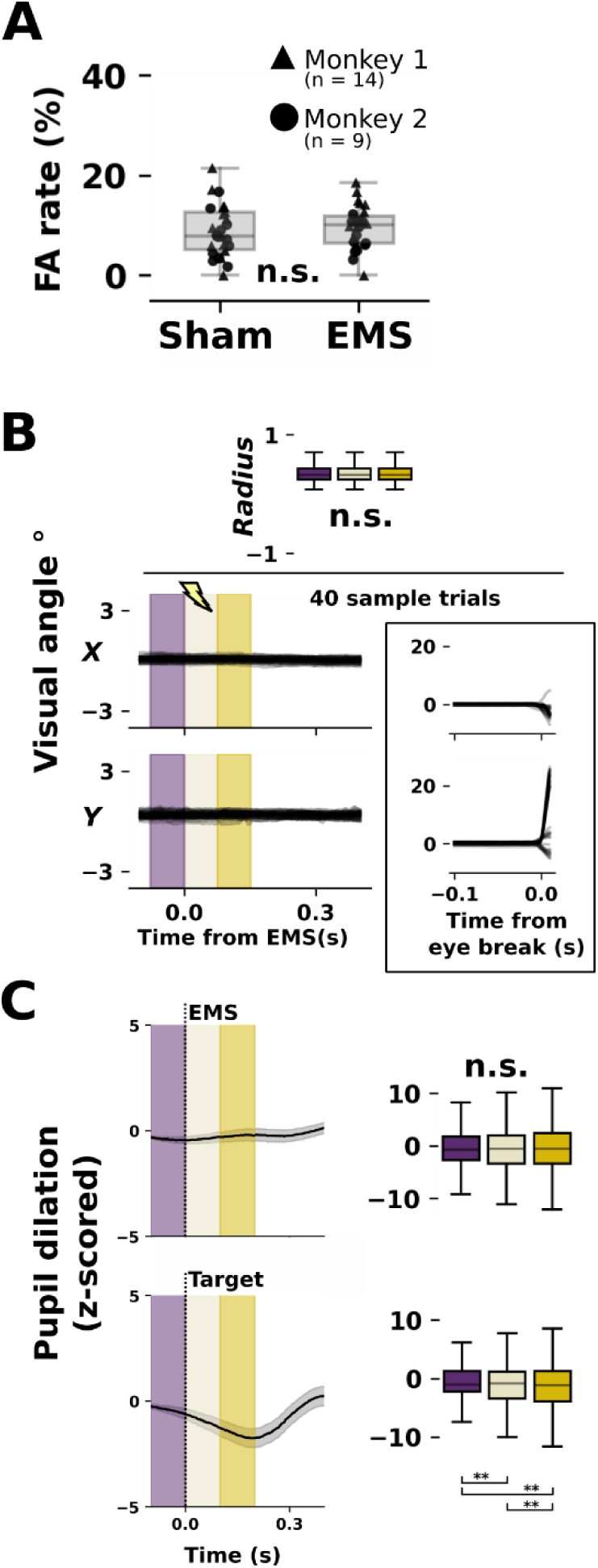
Pulvinar EMS did not elicit eye-movements or false alarms. A) pulvinar electrical microstimulation did not induce lever releases. Boxplots show false alarms in the sham and EMS conditions across sessions. B) no eye movements were elicited following pulvinar EMS. Top: boxplots from sample session of all EMS trials calculated for pre-stimulation (purple) during (tan) and after EMS (yellow). Y-axis denotes the radius of eye position from the center of the screen in degrees of visual angle. Bottom, left: sample 40 EMS trials for X (top) and Y (bottom) axes on the screen. Inset: for comparison, the X and Y positions from 40 sample eye break trials where the animals broke fixation before finishing a trial. C) Similar to panel B, pupil dilation was not affected by EMS (top). For reference, the pupil area was smaller following target presentation (bottom). Due to large pupil dilation drifts, waveforms are normalized for each trial (z-score) according to a −0.2 to −0.1s window prior to time-locking event. Shaded regions denote the standard error of the mean. Boxplot conventions follow Fig. 1.

